# Autocorrelation-based method to identify disordered rhythm in Parkinson’s disease tasks: a novel approach applicable to multimodal devices

**DOI:** 10.1101/2020.08.19.256982

**Authors:** Kenichiro Sato, Tatsuo Mano, Atsushi Iwata, Tatsushi Toda

**Author notes:** Corresponding author (K.S), (A.I). **Data availability:** The data used in this study is publicly distributed from the following websites (http://www.cs.toronto.edu/~taati/index.htm and https://zenodo.org/record/54551#.Xq2oYy3AOql).

## Abstract

**Objective:** We aim to propose a novel method of evaluating the degree of rhythmic irregularity during repetitive tasks in Parkinson’s disease (PD) by using autocorrelation to extract serial perturbation in the periodicity of body part movements as recorded by objective devices.

**Methods:** We used publicly distributed sequential joint movement data recorded during a leg agility task or pronation-supination task. The sequences of body part trajectory were processed to extract their short-time autocorrelation (STACF) matrices; the sequences of single task conducted by participants were then divided into two clusters according to their similarity in terms of their STACF representation. The Unified Parkinson’s Disease Rating Scale sub-score rated for each task was compared with cluster membership to obtain the area under the curve (AUC) to evaluate the discrimination performance of the clustering. We compared the AUC with those obtained from the clustering of the raw sequence or short-time Fourier transform (STFT).

**Results:** In classifying the pose estimator-based trajectory data of the knee during the leg agility task, the AUC was the highest when the STACF sequence was used for clustering instead of other types of sequences with up to 0.815, being comparable to the results reported in the original analysis of the data using an approach different from ours. In addition, in classifying another dataset of accelerometer-based trajectory data of the wrist during a pronation-supination task, the AUC was again highest up to 0.785 when clustering was performed using the STACF rather than other types of sequence.

**Conclusion:** Our autocorrelation-based method achieved a fair performance in detecting sequences with irregular rhythm, suggesting that it might be used as another evaluation strategy that is potentially widely applicable to qualify the disordered rhythm of PD regardless of the kinds of task or the modality of devices, although further refinement is needed.

## Introduction

In the diagnosis and follow-up of patients with Parkinson’s disease (PD), physicians or neurologists examine patients for several signs and symptoms, e.g., bradykinesia, tremor, small steps, posture instability, and freezing of gait (FOG) [1]. In particular, the degree and amount of rhythm irregularity during repetitive tasks (e.g., finger tapping or leg agility) are some of the most frequently evaluated phenomena. The evaluation of these tasks is practically dependent on the neurological assessment performed by physicians or neurologists that can be semi-quantified as scores on the Unified Parkinson’s Disease Rating Scale (UPDRS) [1], which may lead to the limited inter-rater variability [2] for each evaluation item.

A growing body of literature shows that these examinations for PD can be objectively measured using specific devices, such as an accelerometer or motion capture system [2–4]. For example, accelerometers have been used to assess finger tapping [5, 6], pronation-supination [7], tremor [6], gait [8, 9], postural instability [10], FOG [11], and levodopa-induced dyskinesia [12]. In addition, features of gait in PD have been intensively investigated using 3D motion capture in the field of gait analysis [4]. Furthermore, video-based assessment via deep-learning based pose estimators such as the Convolutional Pose Machine (CPM) [13] or OpenPose [14] has been applied to UPDRS tasks as a substitute for conventional 3D motion capture systems, revealing fair prediction performance [15]. Many of these earlier studies employ an evaluation scheme to extract static features from movement sequences with which to discriminate or quantify the degree of abnormal findings.

The general concerns in such evaluation schemes include the complexity of preprocessing procedures and the limited applicability of evaluation algorithms to other tasks/devices. In each algorithmic evaluation, extracting static features (e.g., mean, median, standard deviation, maximum and minimum values) from trajectory sequences or frequency analysis using Fourier transform (FFT) have been predominantly used so far, which requires researchers to standardize all included samples strictly and extract many static features as variables to impute into the discrimination model. In addition, these requirements of the preprocessing procedure make the obtained algorithm device-/task-specific, which results in limited applicability to the different kinds of tasks or to the data obtained using different modality devices.

As one of the solutions to address these points, in this study we attempt to employ autocorrelation (ACF), which we had introduced in our previous study to quantify FOG based on videos of the gait of patients with PD recorded in the hospital hallway [16]. Since the ACF extracts the periodicity representation of the original sequence, it does not require strict standardization of the amplitude of the original sequence. Since semi-quantitative scores evaluating repetitive tasks (e.g., those included in the UPDRS such as finger tapping, leg agility, or toe tapping) are measured based on the regularity of tapping rhythm/speed/amplitude [1], it would be suitable to apply ACF to detect irregularities during the repetitive task, which we aim to propose as a novel method. We then conducted unsupervised clustering for the ACF sequences derived from these raw sequences, which is another feature of our approach. This allows us to identify repetitive sequences with a similar degree of alteration in periodicity during each task, in a data-driven manner, without the need for a complex preprocessing procedure. In addition, there is no need to determine task/device-specific cut-off values.

These strengths might allow this method to be applied to a wide range of repetitive PD tasks evaluating the irregularity of movement, regardless of the modality of devices. To validate our ideas, we used two different datasets obtained from devices of different modalities recording different PD tasks that have been publicly distributed by Li et al. [15] and Piro et al. [7]; the former comprised of a series of estimated 2D coordinates of body joint trajectory during a leg agility task [15] and the latter is an accelerometry-derived sequence during a pronation-supination task [7].

## Methods

### General processing procedure

This is a retrospective study using publicly distributed data. All data handling and statistical analyses were performed using R software (version 3.5.1). Suppose we have the original trajectory sequence of a body part during the tasks recorded at any frequency per second (fps) as in Fig 1A[a]. This ‘raw’ sequence is derived from the serial change in coordinates along with the actual movement of body part during the task, such as the serial change in Y-axis coordinate of ipsilateral knee during the leg tapping task as illustrated in Fig 1A (here we used a free illustration distributed from Irasutoya: https://www.irasutoya.com). This raw sequence was further processed as outlined in Fig 1A; first, the “difference sequence” (Fig 1A[b]) was obtained as the difference series of the raw sequence, which corresponds to the serial changes in the velocity of the sequence (Fig 1A[a]). Next, we applied a sliding window with a certain length and with a certain shift length to these sequences. For the sequence obtained from each sliding window, we applied the FFT (Fig 1A[c]) and autocorrelation (ACF) (Fig 1A[d]) functions, thereby obtaining the resultant vectors. The cumulated vectors along with the sliding form matrices, referred to as short-time FFT (STFT) (Fig 1A[e]) and STACF (Fig 1A[g]) matrices, respectively. For the convenience of calculation that follows later, we took the average of each row of the matrix to derive the averaged STFT (Fig 1A[f]) and STACF (Fig 1A[h]). The difference sequence (Fig 1A[b]) was also used to derive the averaged STFT and STACF.

**Fig 1.**
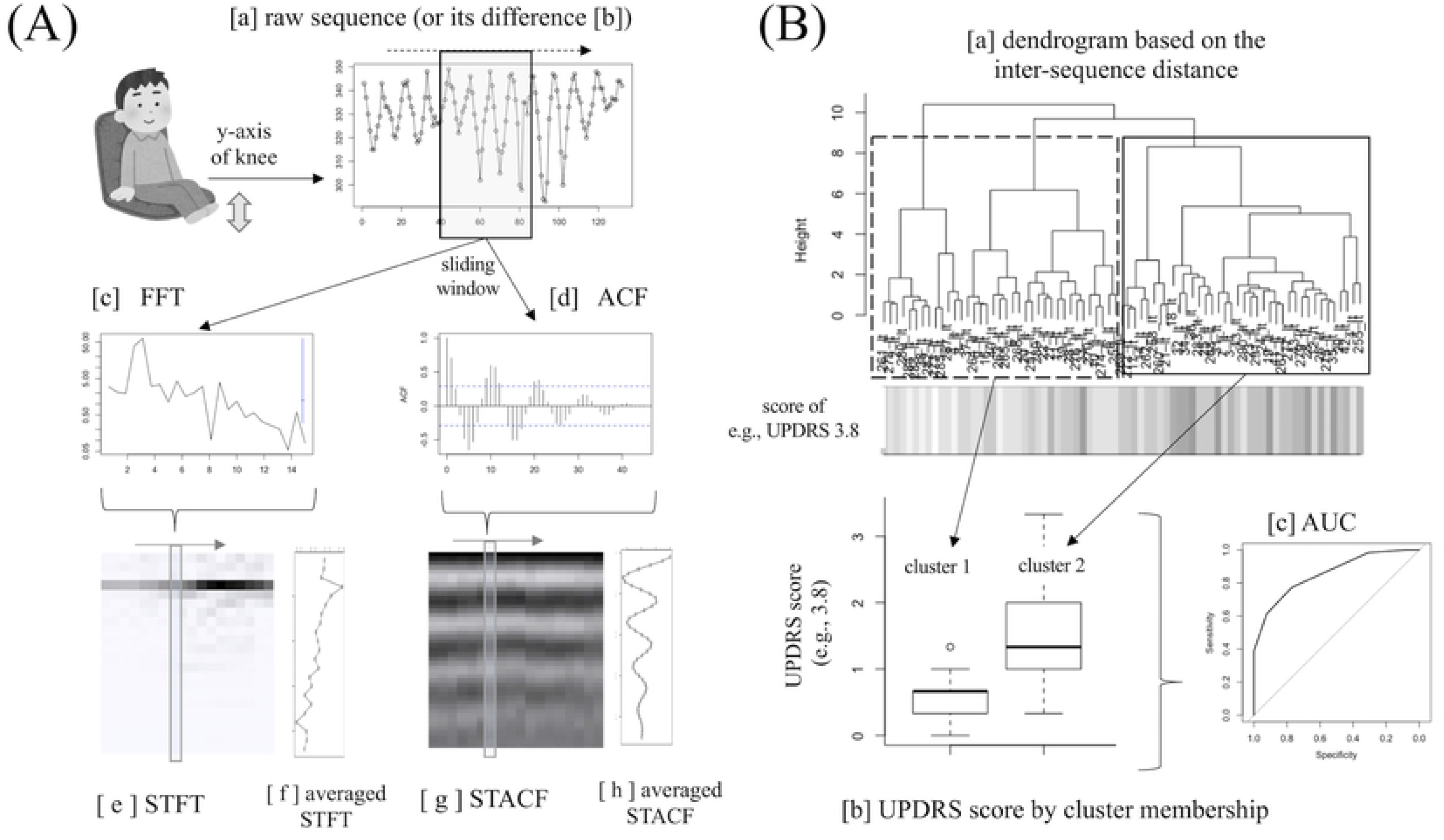
Processing workflow. Fig 1A shows how to derive STFT and STACF sequences from the raw sequence representing the serial coordinate change, and Fig 1B demonstrates how to use the derived six types of sequences for classification of rhythm irregularity, via the clustering of the same type of sequences.

The STACF [17, 18] is an applied form of the STFT [19, 20] and was a method we had proposed previously to apply to the estimated trajectory data of gait in order to quantify FOG [21] from the gait movies of patients with PD recorded in the hospital hallway [16]. The STACF is better than the STFT in terms of frequency resolution within the frequency range of gait: although the STFT is the major method for pitch detection/analysis in acoustic analysis, since the frequency resolution in STFT is given by the reciprocal of the window width (s). Because the window width is several seconds in the tapping movies recorded in actual clinical settings, the frequency resolution in STFT should be lower than that in STACF at any rate used here. The STACF also has the advantage that it can measure and visualize the real-time subtle change in gait rhythms during walking [16].

This produced a set of six sequential vectors from each original raw sequence as follows: original “raw” sequence [a], its “difference” sequence [b], the averaged STFT sequence of the “raw” sequence [f], the averaged STACF sequence of the “raw” sequence [h], the averaged STFT sequence of the “difference” sequence, and the averaged STACF sequence of the “difference” sequence in Fig 1A. We then assessed which of these sequence types may have the strongest association with the degree of rhythmic irregularity in the task, which refers to the semi-quantified UPDRS score that had been provided by the authors of the original data. We conducted sequence clustering using the R package *TSclust* [22] for each of the six sequence types independently (Fig 1B). For example, we have the sequences of a single type from all original “raw” sequences and the inter-sequence distances within the same sequence type were measured and then separated into two clusters according to the dendrogram (Fig 1B[a]) (here we set the number of clusters as two for the purposes of binary classification). The derived dendrogram and task irregularity rating (as determined based on UPDRS score) of each sequence is simultaneously presented in Fig 1B[a] using the R package *WGCNA* [23]. The summary distribution of UPDRS item’s score for each sequence belonging to each cluster (cluster [1] and cluster [2]) is shown in Fig 1B[b], which shows a clear difference in the UPDRS score distribution between cluster [1] and [2], meaning that based on the sequence clustering results we can conversely estimate whether the sequence of interest may have higher or lower UPDRS score regardless of the cut-off value of the UPDRS item of interest. The performance was measured using the AUC (Fig 1B[c]) using the R package *pROC* [24].

To calculate the AUC, we used various distance measures (dynamic time warping (DTW), autocorrelation-based method, Euclidean distances, linear predictive coding ARIMA method, and model-based ARMA method; for details, see [22]) and various clustering methods (“complete,” “ward.D,” and “ward.D2” methods in R [25]). These methods were selected because they are available even if the sequence length differs between the pair to measure the distance. Eventually, for each of the six sequence types, we calculated 15 AUC results according to the 5 (distance measures) × 3 (clustering methods) = 15 combinations of distance and clustering methods. The derived AUC results under a certain distance/clustering method condition were compared using DeLong’s test [24] between the pair of different sequences (as in Fig 2, the lower row), as follows: between the raw difference and STACF sequences, between the STACF and STFT sequences, between the “difference sequence” and the STACF of the difference sequence, and between the STACF and STFT of the difference sequence. These pairs of comparisons were specifically selected in order to examine whether using the STACF as a sequence type in clustering may lead to a better discrimination performance, as well as to restrict the number of pairs to conduct as few statistical tests as possible. The P-value was adjusted using Benjamini-Hochberg method [26] for multiple comparison within these four pairs of comparison, and a false discovery rate (FDR)-adjusted p-value less than 0.05 was regarded as statistically significant.

**Fig 2.**
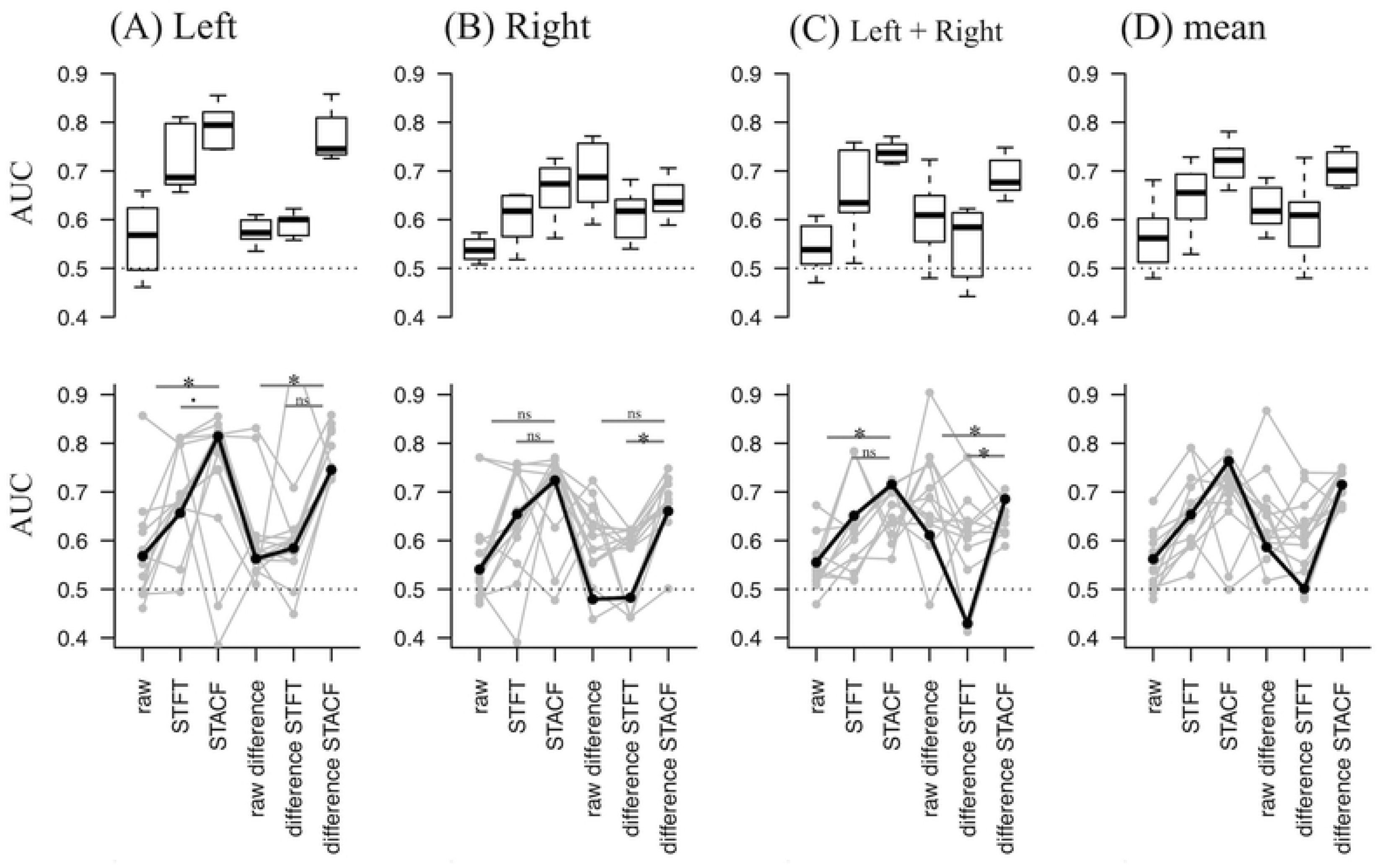
Performance results from dataset (1) In the upper row, derived AUC distribution across 15 distance/clustering method conditions for each of six sequence types (for columns from left to right). Sequences of different laterality were separately analyzed (i.e., left leg, right leg, and both legs) according to the original article of this data. In the lower row, AUC results of one specific distance/clustering condition are plotted, and the results of statistical tests in AUC comparison are shown on the presented pairs.

### Dataset (1): estimated joint trajectory during a leg agility task

We downloaded a publicly available dataset from http://www.cs.toronto.edu/~taati/index.htm in March 2020, which included data from the study by Li et al. [Li 2018]. The data consisted of 2D body joint trajectories (serial X and Y-axis coordinates) from task videos as estimated using Convolutional Pose Machines (CPM) [13], a deep learning-based pose estimation tool to estimate the position of human body parts/joints including the face, right/left-sided shoulder, elbow, wrist, hip, knee, and ankle in the picture (or each frame of the movie). The obtained data are the time series vectors of these joint coordinates recorded at a frequency of 30 frames per second (fps) during each specific task of communication, drinking, leg agility, and toe tapping. Among them, we used the Y-axis sequential data of the knee during the leg agility task (UPDRS 3.8 [1]) from the ipsilateral knee, where the participants raise and stomp either the left or right foot while sitting on a chair. In the original article, this sequence has already been filtered by low-pass filter with a cut-off of 5 Hz. The UPDRS 3.8 score in each leg agility task evaluated by neurologists (average of two or more neurologists’ evaluation) is annotated for each sequence. Sequences in which UPDRS 3.8 evaluation was not available were excluded from the analysis. This “raw” sequence (the Y-axis sequence of the moving knee during the leg agility task) is further processed as outlined in Fig 1A. In obtaining the STFT and STACF, we applied a sliding window that was 45 frames wide (= 1.5 seconds) with a shift length of 5 frames (= 0.167 seconds), which were determined mainly based on the length of each sequence ranging from 57 to 329 frames. The performance of binary classification achieved in the original study [15] was AUC = 0.842 and 0.699 for the average of sequences from the left and right legs, respectively.

### Dataset (2): accelerometer-derived data during the pronation-supination task

Next, for further validation of our proposed method, we applied our method to another dataset from different UPDRS tasks using different device modality. We downloaded publicly distributed data from https://zenodo.org/record/54551#.Xq2oYy3AOql in April 2020, which were originally used in the study reported by Piro et al. [7]. This data is consisted of time series data of 3-axis acceleration (x, y, and z) measured using an accelerometer attached to the participants’ wrist during a pronation-supination task, as determined using UPDRS 3.6 [1] where the participant lifted one arm into a horizontal position and turned the palm up and down 10 times while sitting on a chair. The UPDRS 3.6 score of each pronation-supination task evaluated by neurologist(s) is annotated for each sequence.

We first normalized each of the 3-axis (x, y, and z) acceleration sequences within which the amplitude ranged 0–1, and then obtained the synthetic acceleration sequence from these 3-axis acceleration sequences. This sequence was normalized and further centralized by subtracting the median value of the sliding window (20 frames), making the sequence’s baseline closer to 0. The obtained is the sequence in Fig 3A[a]. Since this sequence is the first-derivative of the velocity sequence and the velocity sequence is also the first-derivative of the trajectory sequence, the sequence in Fig 3A[a] was integrated twice. We used the calculated sequence (Fig 3A[c]) as the “raw” sequence as in dataset (1). In applying the STFT and STACF, the window width was set to 100 frames and the shift length was set to 10. In this dataset (2), the AUC had not been reported in the original article.

**Fig 3.**
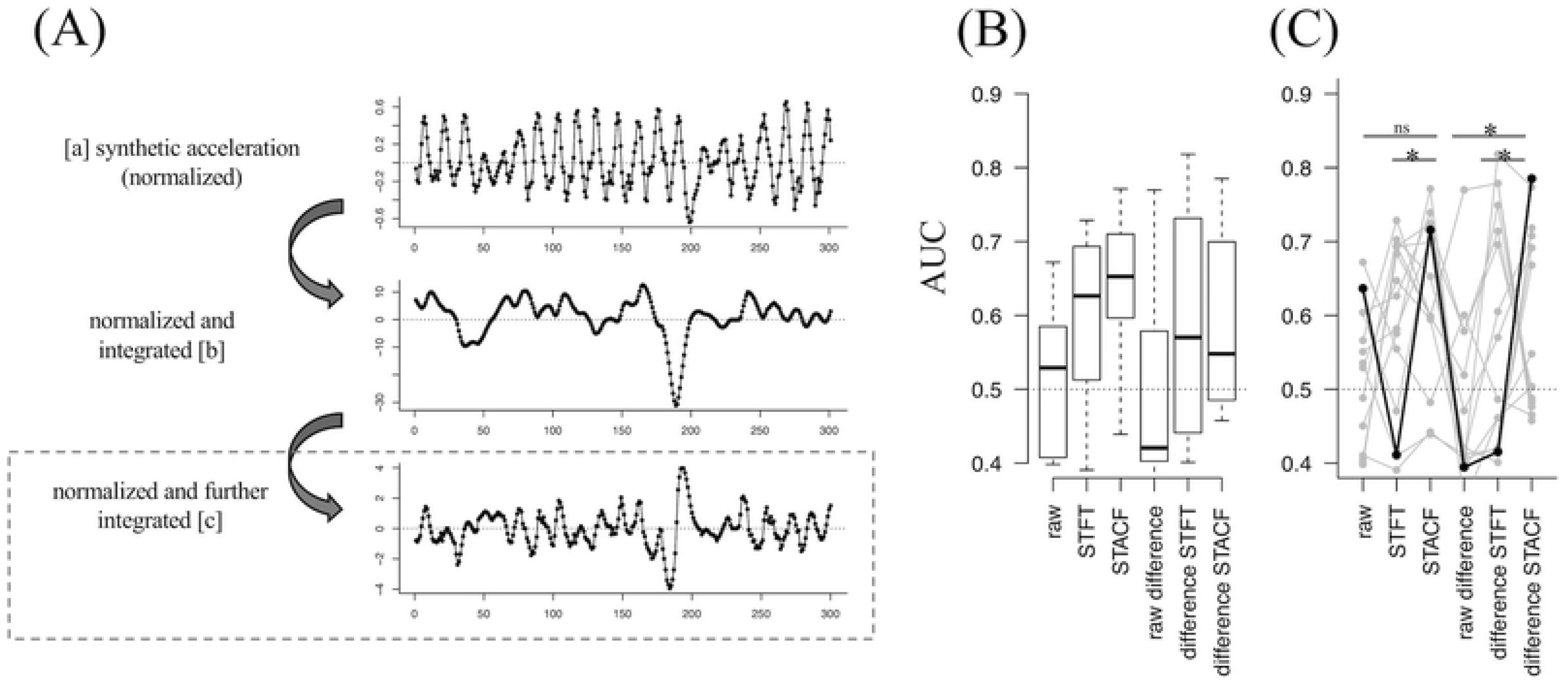
Procedure and the performance results from dataset (2) Fig 3A shows the preprocessing procedure for accelerometer-derived serial data as in Fig 1A so that we can apply the same clustering procedure as of dataset (1). Fig 3B & Fig 3C are the result for this dataset (2), showing similar tendency as of dataset (1).

### Ethics

This study was approved by the University of Tokyo Graduate School of Medicine institutional ethics committee [ID: 2339-(3)]. Informed consent was not required for this study. The work was conducted in accordance with the ethical standards laid out in the Declaration of Helsinki, 1964.

## Results

### Performance results: dataset (1)

In total, we included 150 sequence samples from dataset (1), which consisted of left- and right-sided sequences obtained from 75 unique trials conducted by nine participants. The AUC performance results are summarized in Fig 2 (boxplot in the upper row and superimposed plot in the lower row), where the AUC distribution across 15 distance/clustering method conditions for each of the six sequence types (for columns from left to right, original “raw” sequence and its STFT and STACF, and the “difference” sequence and its STFT and STACF). Sequences of different laterality were separately analyzed as in the original study [15], i.e., results using sequences from left legs (Fig 2A), right legs (Fig 2B), and sequences from either left or right legs (Fig 2C). Fig 2D is the geometric mean of left-sided and right-sided results. Briefly, results based on the STACF sequence appeared to show a better performance distribution than those of the other types of sequences (Fig 2, upper row).

When we arbitrarily focused on a specific condition to examine the detailed performance of the STACF sequence (as highlighted with solid [not gray] lines on the lower row of Fig 2; DTW as distance method and ‘ward. D’ as clustering method), the AUC = 0.815 of the STACF sequence was the highest among the sequences from the left leg (Fig 2A, lower row) and was significantly better than the AUC = 0.568 of the “raw sequence” (FDR = 0.021 in Delong’s test, marked with * in Fig 2A). In the same condition, the AUC = 0.715 for STACF was the highest among the sequences from the right leg (Fig 2B, lower row) but was not significantly better than the AUC of the raw or STFT sequences. Furthermore, the AUC = 0.724 of STACF was also the highest among the sequences from either the left or right legs (Fig 2C, lower row) and was significantly better than the AUC = 0.541 of the “raw” sequence (FDR = 0.031, marked with * in Fig 2C).

These results suggest that clustering of the STACF sequence in a specific condition might lead to statistically better discrimination performance than clustering of the raw sequence (or its difference sequence).

### Performance results: dataset (2)

Next, for further validation of our proposed method, we applied our method to another dataset (2). In total, we included 101 sequence samples, which consisted of left- and right-sided sequences obtained from 26 unique participants. The AUC performance results are summarized in Fig 3 (boxplot in Fig 3B, and superimposed plot in Fig 3C). Sequences with different laterality were not separately analyzed in the original study [7]. The results showed a similar performance distribution as in dataset (1), where the STACF-based results appeared to show a better performance distribution than that of the other types of sequences (Fig 3B).

Next, we focused on the same conditions as in dataset (1), DTW as a distance method and “ward. D” as a clustering method; the STACF sequence (AUC = 0.715) had significantly higher AUC than the “STFT” sequence (FDR < 0.001, marked with * in Fig 3C). The STACF of the difference sequence (AUC = 0.785) had a significantly higher AUC than the “raw” or “STFT” sequences (both FDR < 0.001, marked with * in Fig 3C).

These results suggest that the better discrimination performance when clustering STACF rather than the raw or STFT sequences is preserved for the different types of sequences with different tasks and different modalities.

## Discussion

Neurological findings from patients suspected of having PD are seldomly measured using objective devices such as an accelerometer or motion capture system [2–4], probably due to the barriers to their use in daily practice; these devices are time-, labor-, space-, and cost-consuming. As a result, the evaluation of each examination task is currently solely dependent on the neurological assessment of physicians or neurologists, that may be semi-quantified as scores on the UPDRS [1]. This may lead to the limited inter-rater variability [2] in each evaluation item. As one of the options for achieving objective evaluation, video-based assessment via deep learning-based pose estimators such as the CPM [13] or OpenPose [14] is available; these pose estimators automatically estimate the joint coordinates of the person in the pictures or videos obtained using a monocular camera without requiring external scales or markers, and therefore can be used as a substitute for a conventional 3D motion capture systems. Indeed, when assessing the motor symptoms of PD, the degree of disability in some UPDRS tasks and degree of levodopa-induced dyskinesia recorded in movies has been shown to be measurable using a combination of pose estimation and machine learning [15]. Furthermore, we previously reported a study that applied a pose estimator to videos of the gait of patients with PD recorded in the hospital hallway [16] and proposed a novel method using STACF to quantify FOG [21].

In line with these earlier studies that applied pose estimators for vision-based assessment of PD symptoms, in this study, we aimed to propose a novel method to evaluate the degree of disordered rhythm in repetitive tasks in PD patients. As expected, the results using the STACF sequence for clustering generally showed a better discriminant performance than using other sequences such as the raw sequence or STFT sequence. By choosing the appropriate condition for the sequence clustering, we could obtain the AUC performance results that are compatible to those reported in the original study [15], where the average AUC was 0.842 and 0.699 for sequences from the left and right legs, respectively. In addition, even when the current method was applied to another dataset from a different task and different modality, the best discrimination performance was also observed when clustering the STACF sequence. These results suggest that our proposed method of clustering the STACF sequence may be widely applicable as an alternative evaluation scheme to conventional approaches for the objective assessment of disordered rhythms in PD patients.

The potential of the wide applicability of our method is further explained by the following points. The STACF extracts temporal changes in frequency within a short time window. Our method does not require the scale of all included sequences to be strictly standardized. This means the relative position between the camera and participants during the same task does not always need to be fixed for all recordings, and in some case zooming in/out during recordings is permitted; such robustness against recording conditions would increase the capacity to incorporate sequential data estimated using a pose estimator [15, 16], derived from the videos recorded in different recording environments or clinical situations. Moreover, our proposed method can also be applied to the tapping task of other body parts, such as finger tapping or toe tapping. Currently, since the CPM library is not equipped with the function to estimate finger or toe trajectories, we need additional pose estimator libraries to extract the trajectory of these tasks.

In addition, by applying ACF to each sliding window but not to the whole sequence, we could unify the length of the obtained averaged STACF sequences across all sample sequences, even if the length of original raw sequences differed greatly. Although we used sequence clustering to perform a binary classification in this study, we also can impute the STACF as the temporal representation of the raw sequences for each sample into a machine learning scheme, where we can expect further improvement in the predictive performance with further investigation.

As the method of measuring the distance between sequences, we focused on the DTW because it achieved a good discrimination performance. The DTW is a time-series clustering algorithm that calculates the distance between two waveforms’ sequential patterns (= 0 between the same sequences) and has an advantage that it can compare two waveform data of which the length or phases differ significantly. Since the tapping sequences diagnosed as “normal” but with different tapping speeds result in STACF sequences with similar waveforms but mildly different phases (varying lag peaks), the use of the DTW is appropriate to measure distances in sequence clustering.

Our study has some limitations. First, our proposed methods require further external validation by incorporating additional sequence data from different tasks obtained using different modalities in different environments or facilities. Second, it is uncertain why the basic performance in dataset (1) differs between the sequences from the left and right legs, although this may be due to the characteristics of the original data. Third, the strength of STACF focusing on the periodicity of sequence may in turn impair sensitivity to severely impaired tapping that leads to persistently lowered amplitude of tapping/pronation-supination due to the genuine presentation of severe parkinsonism. Fourth, there are some technical concerns such as whether the number of clusters can be extended (from two in this study), that we should further explore the appropriate width and shift length of the sliding window, that not all possible types of distance methods and clustering methods are examined, or that averaging the STACF and STFT matrices may overlook temporal characteristics in the change in periodicity, especially in longer sequences. In addition, the hierarchical clustering we used in this study may not work well for sequential data that is too long, e.g., more than thousands of time-steps, so our method may not be applicable to the long time series data such as that obtained from monitoring patients’ disease status based on their daily activity.

To conclude, we conducted clustering for the STACF-derived from trajectory sequences, which allowed us to identify task samples with a similar degree of alteration in periodicity during each task. Because our approach is a data-driven classification method that does not require complex preprocessing procedure nor strict standardization across samples, our proposed method may be used as another evaluation scheme that is widely applicable to any PD tasks (including finger tapping and toe tapping) to evaluate irregularity of movement, regardless of the modality of the measurement devices.

## Acknowledgments

This study used the data distributed by Li, et al (http://www.cs.toronto.edu/~taati/index.htm) and Piro, et al (https://zenodo.org/record/54551#.Xq2oYy3AOql), which the authors have accessed on April 2020.

